# Global survey of secondary metabolism in *Aspergillus niger* via activation of specific transcription factors

**DOI:** 10.1101/2024.07.18.604165

**Authors:** Cameron Semper, Thi Thanh My Pham, Shane Ram, Sylvester Palys, Gregory Evdokias, Jean-Paul Ouedraogo, Marie-Claude Moisan, Nicholas Geoffrion, Ian Reid, Marcos Di Falco, Zachary Bailey, Adrian Tsang, Isabelle Benoit-Gelber, Alexei Savchenko

**Affiliations:** Department of Microbiology, Immunology and Infectious Disease, University of Calgary, 3330 Hospital Drive, Calgary, Alberta, T2N 4N1, Canada; Centre for Structural and Functional Genomics, Concordia University, 7141 Rue Sherbrooke Ouest, Montreal, Quebec, H4B 1R6, Canada

**Keywords:** Secondary metabolism, transcriptional regulation, natural products

## Abstract

Genomics analysis confirmed the status of the filamentous fungi as a rich source of novel secondary metabolites; however, the discovery of these compounds is hampered by the cryptic nature of their biosynthetic pathways under laboratory conditions. Consequently, despite substantial research effort over the past decades, much of the secondary metabolome remains uncharacterized in fungal organisms. Our manual curation of biosynthetic gene clusters (BGCs) in the *Aspergillus niger* NRRL3 genome revealed that only 13 of 86 BGCs have had their cognate secondary metabolite products confirmed or reliably inferred. We also identified 60 transcription factors associated with cryptic BGCs. To further characterize *A. niger* secondary metabolism, we created a collection of strains each overexpressing a single BGC-associated transcription factor. We analyzed the strain collection using a standardized pipeline where we monitored phenotypic changes and compound production using mass spectrometry. Strains showing evidence of secondary metabolism activation were selected for gene expression analysis. Our approach resulted in the production of multiple potentially novel secondary metabolites and linked a specific BGC to tensidol production in *A. niger.* More broadly, this study found evidence counter to the existing paradigm of BGC expression controlled by co-localized transcription factors, lending credence to the emerging picture of a complex regulatory network governing fungal secondary metabolism.

**Significance Statement:** Fungi produce an array of chemically diverse compounds that are routinely found to harbour valuable bioactivity. The products of secondary metabolism, these compounds have been a source of antimicrobials, anti-cancer agents, and other biopharmaceutical compounds termed natural products. Despite their demonstrated economic value, much is still unknown about the biosynthesis, regulation, and identities of these compounds. This study adopted a genome-wide approach to improve our understanding of the regulatory mechanisms that control fungal secondary metabolism, improving our ability to investigate the pathways responsible for natural product production.

## Introduction

Fungal secondary metabolites are organic compounds that are rich in chemical diversity and include many molecules of clinical and industrial significance (1). Examples include anti-microbials (e.g. penicillin, echinocandins), anti-cancer compounds (e.g. aurantiamine, wortmannin), immunosuppressants (e.g. cyclosporin, mycophenolic acid) and anti-cholesterolemic compounds (e.g. statins) (2–4). Filamentous fungi have traditionally been a rich source of secondary metabolites and display a capacity to produce compounds that are both beneficial through exploitation (e.g. gibberellic acid) and harmful (e.g. aflatoxins, ochratoxin) to humans (5, 6). The availability of fungal genome sequences, combined with improved bioinformatic tools, has further highlighted the wealth of untapped biosynthetic potential of filamentous fungi for the production of novel secondary metabolites (7). Despite the clinical, industrial, and economic potential associated with fungal secondary metabolism, the majority of the secondary metabolome of these organisms remain poorly characterized. As a result, substantial effort has been made towards improving our understanding of the genetics and processes that control expression of these small molecules with an eye towards combatting spoilage due to mycotoxin production, discovering new bioactive molecules, and linking known metabolites to their corresponding biosynthetic enzymes.

The biosynthetic genes responsible for production of secondary metabolites are typically co-localized into discrete biosynthetic gene clusters (BGC) within fungal genomes (8). BGCs contain two or more genes that contribute to the biosynthesis of a secondary metabolite and can be defined by the enzyme class of the corresponding backbone enzyme(s) (9). Backbone enzymes catalyze extension of the polymer chain and include polyketide synthases (PKS), non-ribosomal peptide synthetases (NRPS), hybrid PKS/NRPS, terpene synthases, fatty acid synthases (FAS) and dimethylallyl tryptophan synthases (DMAT). BGC may also encode for proteins involved in decorating the backbone through various enzymatic activities (e.g. redox, methylation), transporters or efflux pumps, and in some cases transcription factors (TF) that can play a role in regulating the overall biosynthetic pathway (8). Based on the class of backbone enzyme present within a BGC, general predictions can made about the chemistry of the cognate secondary metabolite(s); however, predicting the final chemical structure and corresponding biological function of the metabolites produced by BGCs remains elusive necessitating empirical characterization. Further hampering the characterization and exploitation of the fungal secondary metabolome is the fact that the majority of BGCs found in typical ascomycete species are transcriptionally silent under laboratory conditions (10).

Spurred by advances in molecular biology, considerable effort has been invested into awakening silent BGCs to elicit production of their corresponding secondary metabolites. Approaches include modification of culture conditions (pH, temperature), co-culturing with other organisms (11, 12), deleting and/or overexpressing master or co-localized regulators (13), heterologous expression of BGCs (14) or *in vitro* reconstitution of entire biosynthetic pathways (15). While each of these techniques has been successful for identifying a small number of novel secondary metabolites, they are primarily used to parse only a subset of BGCs in a given organism. In cases where novel metabolites are discovered, they are typically produced in quantities insufficient for structural characterization or downstream functional characterization thus necessitating further optimization.

*Aspergillus niger* is a haploid filamentous fungus that has been used extensively for the production of organic acids (e.g. citric acid and gluconic acid) and carbohydrate degrading enzymes (e.g. glucoamylase) (16). It is also used as a model organism for the study of fungal systems, with comprehensive genetic tools available including a recently established protocol for gene editing using CRISPR/Cas9 (17, 18). Publicly available genome sequences of *A. niger*, including the manually curated NRRL3 genome, confirmed it to be a rich source of cryptic BGCs (19).

In this study, we report the manual curation of all BGCs within the *A. niger* NRRL3 genome, which included the identification of 60 cryptic BGCs featuring co-localized Zn_2_Cys_6_ TFs. As a tool to advance the study of *A. niger* secondary metabolism, we generated an overexpression strain collection comprised of 58 *A. niger* NRRL3-derived strains, each overexpressing a single Zn_2_Cys_6_ TF. We then analyzed each strain using a standardized workflow that involved phenotypic profiling, untargeted metabolomics and transcriptomics (Fig 1). Previously characterized secondary metabolites from *A. niger* were well-represented within our strain library and their production could be reliably attributed to overexpression of specific TFs. Analysis of our strain library also found a number of unique metabolomic features associated with activation of cryptic BGCs, leading to correlation of a specific BGC to tensidol B production. Gene expression analysis of the strain library revealed multiple instances of cross-reactivity between BGCs and underscored the complex regulatory network that governs fungal secondary metabolism.

**Figure 1.**
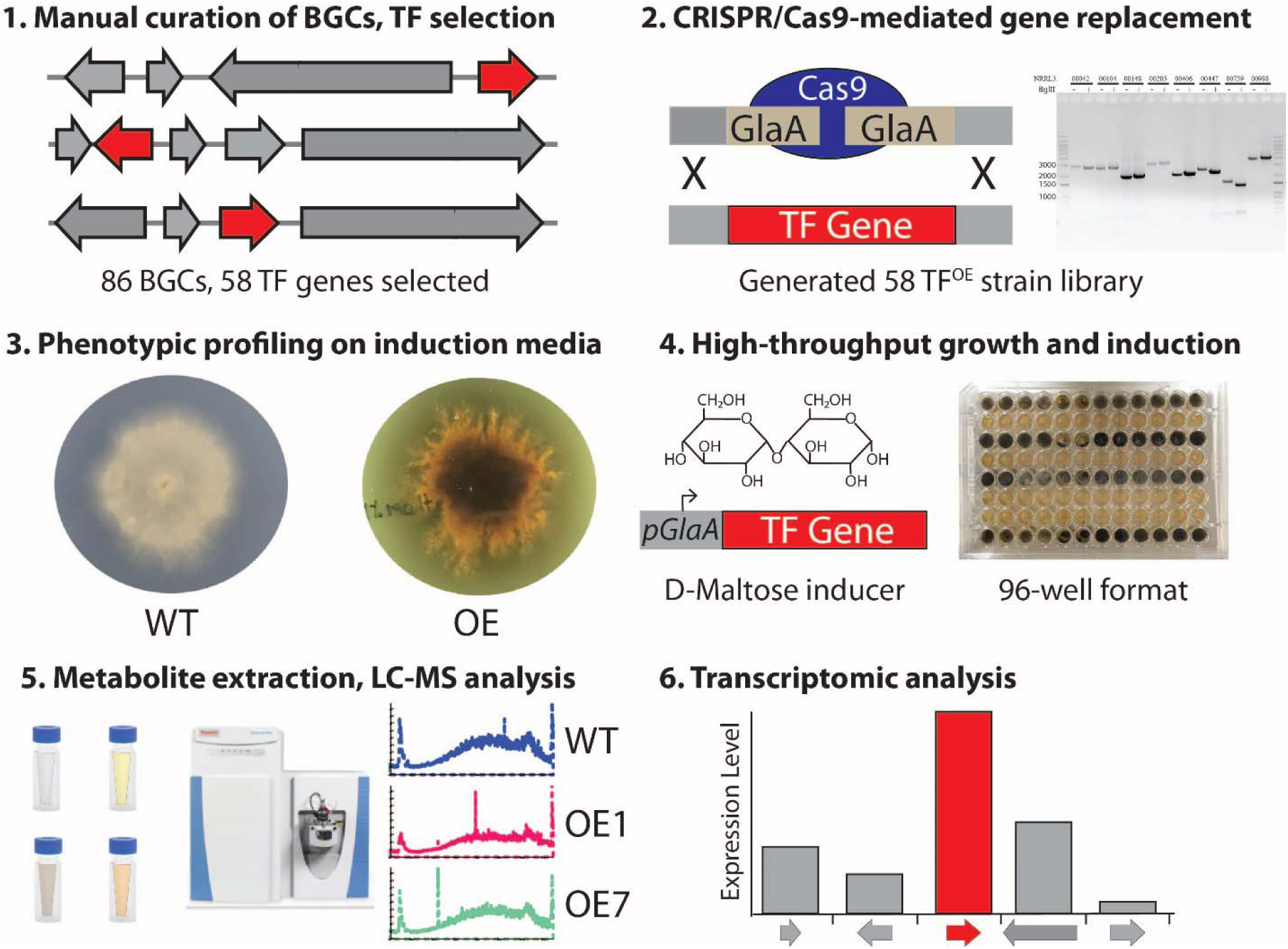
Workflow for investigating secondary metabolism using overexpression strain collection. **1)** the NRRL3 genome was manually curated and all genes localized to BGCs were catalogued. 58 BGC-associated TF genes were identified and selected for overexpression. **2)** using CRISPR/Cas9, each TF was inserted into the glucoamylase locus by replacing the coding region of the *glaA* gene resulting in the generation of a 58-member overexpression strain library. **3)** the strain collection was plated on induction (maltose) media for phenotypic profiling. **4)** strains were cultured in 96-well format on induction media. **5)** metabolite extraction and untargeted LC-MS analysis of samples was conducted. Results were compared to existing *A. niger* metabolite databases to positively identify metabolites where possible. **6)** RNA-seq was performed two-hours post-induction on a subset of strains showing altered metabolite profiles to measure changes in gene expression resulting from overexpression of TF.

## Results

### Manual curation of Biosynthetic Gene Clusters (BGC) in A. niger NRRL3

The publicly available *A. niger* NRRL3 genome sequence contains coverage of all eight chromosomes including 16 telomeric regions and manually verified gene models [17]. We used the bioinformatic tools SMURF (20) and antiSMASH (21, 22) to identify BGCs that may be involved in secondary metabolite synthesis, as well as InterProScan to identify protein domains that correspond to proteins involved in backbone production of secondary metabolites. Results of this curation effort identified a total of 86 BGCs (Table 1), distributed across all eight chromosomes (Fig. 2). Notably, this number of predicted BGC is greater than previously published estimates based on similar analyses of the *A. niger* CBS513.88 genome [18, 19]. Within these 86 BGCs in *A. niger* NRRL3, we identified 99 genes encoding backbone-type enzymes, the majority of which were classified as PKS (n = 40), NRPS (n = 38), or hybrid PKS/NRPS (HPN) (n=9). We also identified a small number of terpene synthase (n=6) and fatty acid synthase (n=5) encoding genes in BGCs. In addition, the 86 BGCs featured 69 genes encoding for Zn_2_Cys_6_ TFs, 53 genes for predicted transporters and more than 300 additional genes encoding predicted tailoring enzymes as well as 77 genes encoding proteins of unknown function. Cross-referencing these BGCs against gene expression data from publicly available gene expression data for *A. niger* CBS513.88 (FungiDB) indicated that most of these BGCs lacked a gene expression profile, consistent with previous results showing >70% of *A. niger* BGCs appear to be transcriptionally silent under laboratory growth conditions [20]. Among BGCs identified in *A. niger* NRRL3, nine have been previously characterized experimentally in terms of production of specific secondary metabolites and two have had their secondary metabolite products inferred on the basis of sequence similarity with characterized BGCs from other organisms (Table 1). Specifically, the NRRL3_02178 – NRRL3_02189 BGC has been proposed to support the production of fumonisin based on similarity to corresponding BGCs in *Fusarium verticillioides* and *Gibberella moniliformis* (23–25), while the expression of NRRL3_07873 – NRRL3_07888 BGC has been predicted to produce malformin A2 and C, based on similarity to a characterised BGC is *A. brasiliensis* (26). We also identified genes homologous to the ones responsible for production of naptho-γ-pyrone and DHN-melanin in *A. fumigatus;* however, these genes are not co-localized into a specific BGC in the *A. niger* NRRL3 genome. In total, our manual curation efforts revealed that only 13 out of 86 BGCs have had their secondary metabolite products confirmed or reliably inferred, suggesting a systematic undertaking to explore the endogenous biosynthetic potential of the remaining 73 clusters is warranted.

**Figure 2.**
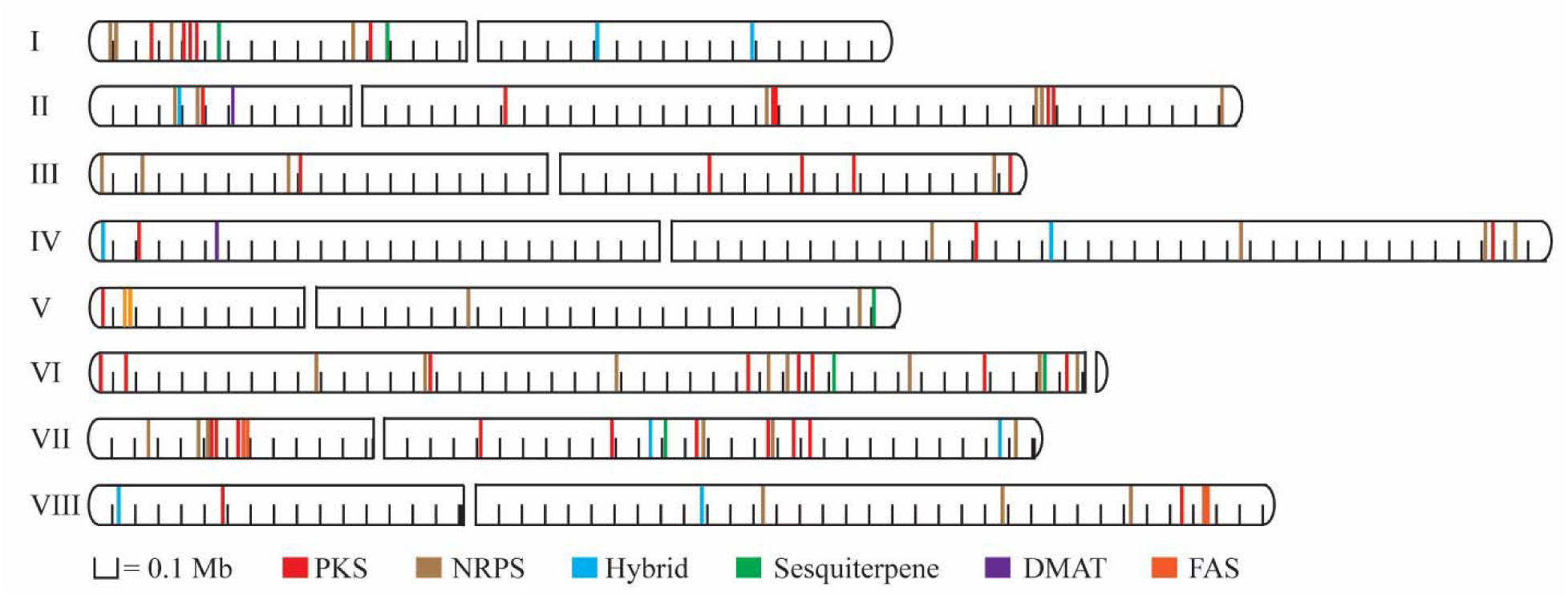
Genomic locations of BGC in *Aspergillus niger* NRRL3. Location of all 86 BGCs shown on chromosome maps, coloured by backbone enzyme-type

**Table 1.**
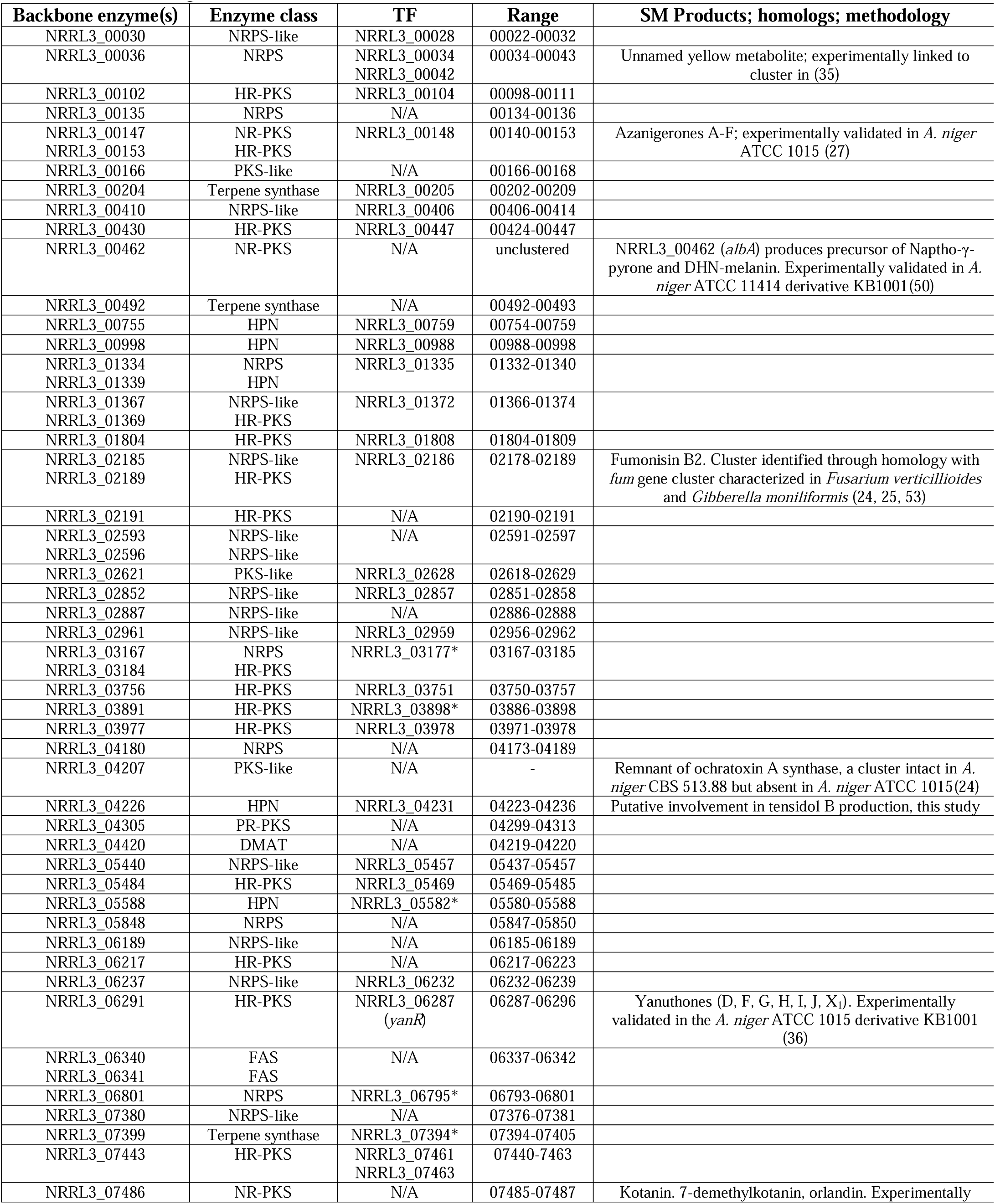
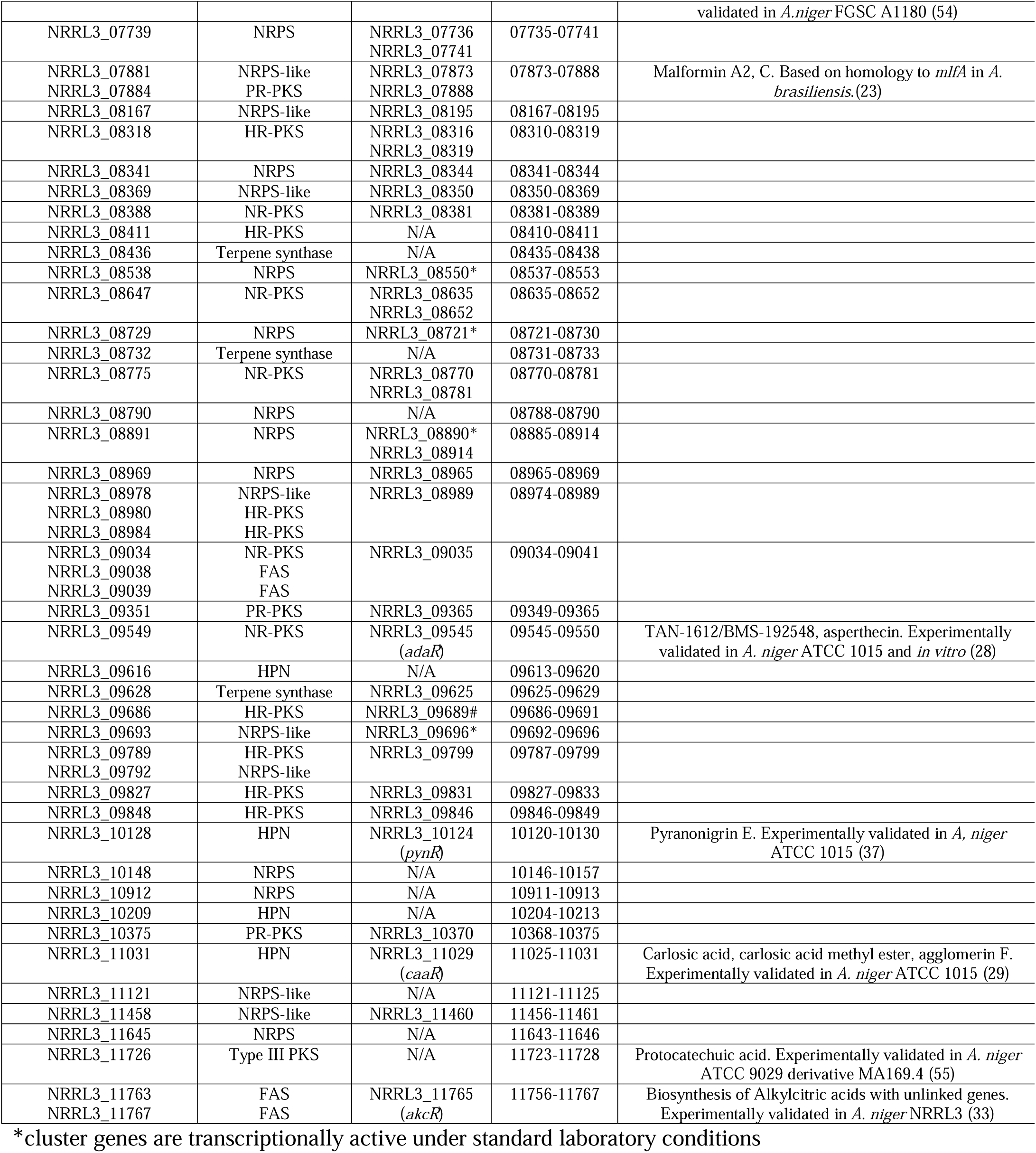
BGC in *A. niger* NRRL3.

### Construction of the A. niger NRRL3 TF overexpression (TF^OE^) strain collection

Previous studies have shown that overexpression of co-localized TF can activate expression of the corresponding BGC in a fungal genome resulting in the production of secondary metabolites (26). This specific strategy has led to characterisation of BGCs in *A. niger* involved in production of azanigerone A-F, agglomerin F and carlosic acid, and TAN-1612 and BMS-192548 (27–29). However, to our knowledge, this strategy has never been applied to investigate a wider spectrum of the *A. niger* NRRL3 secondary metabolome. Accordingly, we decided to pursue the genome-wide overexpression of TFs associated with cryptic BGCs.

As mentioned above, our analysis revealed 69 BGC-associated TF genes, of which 60 appeared to be transcriptionally silent under previously tested growth conditions. The nine TF genes showing expression under previously tested growth conditions were not included in our pipeline for generating overexpression strains and two of the TFs, NRRL3_09689 and NRRL3_11460, were recalcitrant to cloning and after several attempts were omitted as well. The remaining 58 TFs were selected for overexpression, and we undertook construction of the *A. niger* NRRL2270 (ATCC11414) (a spontaneous derivative of ATCC1015 (30)) overexpression strain collection where each of these 58 TF was individually targeted (Fig. 2). To generate the TF^OE^ strains we used *Streptococcus pyogenes* CRISPR/Cas9-mediated integration (17) of each of the 58 TF genes into the glucoamylase A locus, thereby replacing the coding region of the glucoamylase A region with the gene encoding a TF. The integration site selected was downstream of the strong inducible promoter (*P_glaA_*) that has been shown to drive high levels of gene expression during growth on maltose (31, 32). Using this approach we successfully generated 58 TF overexpression strains, with each TF gene integration verified by PCR amplification and restriction enzyme digestion (Figure S1.).

### Phenotypic analysis of overexpression strains

To estimate the potential impact of individual TF gene overexpression on metabolite production we first monitored our *A. niger* NRRL2270 strain collection for potential phenotypic changes. Distinct phenotypic features such as pigmentation (27), delayed or reduced growth (33), or changes in sporulation (34) are often early indicators of changes in fungal secondary metabolism. All 58 TF overexpression strains were grown on transformation media and induction media (2% maltose as carbon source) and monitored over seven days. In at least 14 strains we observed unique phenotypic markers including pigmentation, delayed/slow growth, and impaired sporulation (Figure S2).

Pigmentation during growth on transformation or induction media was observed in 8 of the 14 strains (Figure S2). This group included six strains where the overexpressed TF is associated with previously characterised or functionally predicted BGC, including NRRL3_00042, NRRL3_00148, NRRL3_06287, NRRL3_09545, NRRL3_10124 and NRRL3_11029. We had previously described the NRRL3_00042 overexpression strain as being involved in production of a yellow pigment (35). Overexpression of NRRL3_00148 resulted in production of a distinct yellow-orange pigment on induction media. This TF is associated with a BGC shown to produce several azaphilone compounds including the yellow-coloured azanigerone A compound (27). Yellow pigmentation was also observed in the NRRL3_06287 overexpression strain, which co-localizes with the BGC previously linked to production of yanuthone compounds (36). For this strain, the phenotypic changes were most evident when the strain was grown on transformation media using sucrose as a carbon source but could also be seen when the strain was plated on induction media. One of the most prominent instances of pigment production was in response to overexpression of NRRL3_09545. The BGC associated with this TF produces multiple compounds including TAN-1612, a yellow aromatic polyketide (28). NRRL3_10124 overexpression also produced noticeable yellow pigmentation during plating on induction media. This TF has been linked to a BGC that produced the yellow compound pyranonigrin E (37). Finally, the NRRL3_11029 overexpression strain displayed pink pigmentation during growth on transformation media and at later stages of growth on induction media (29).

Other phenotypic markers that could be used to distinguish overexpression strains were also associated with overexpression of TFs linked to previously characterized BGCs. Overexpression of NRRL3_02186, a TF linked to production of the fumonisin class of mycotoxins, resulted in reduced growth overall and impaired sporulation (Figure S2). The NRRL3_11765 overexpression strain showed irregular branching and impaired sporulation. This TF was previously shown to regulate a BGC linked to the production of alkylcitric acids (33).

In six of the strains where we observed unique phenotypic traits (Figure 3), the corresponding TFs were not associated with any previously characterized BGC. Overexpression of NRRL3_00205 resulted in a phenotype displaying an irregular branching shape and strong black/brown pigmentation (Figure S2). NRRL3_01335 overexpression resulted in distinct changes in sporulation. The NRRL3_04231 overexpression was marked by delayed growth and an irregular branching phenotype. Strong yellow pigmentation was observed in response to overexpression of NRRL3_09035 (Figure S2). Finally, the TF^OE^ strains overexpressing NRRL3_07741 and NRRL3_10370 displayed an accelerated growth rate compared to the parental strain (NRRL2270), which was accentuated in later stages of culture (Figure S2).

**Figure 3.**
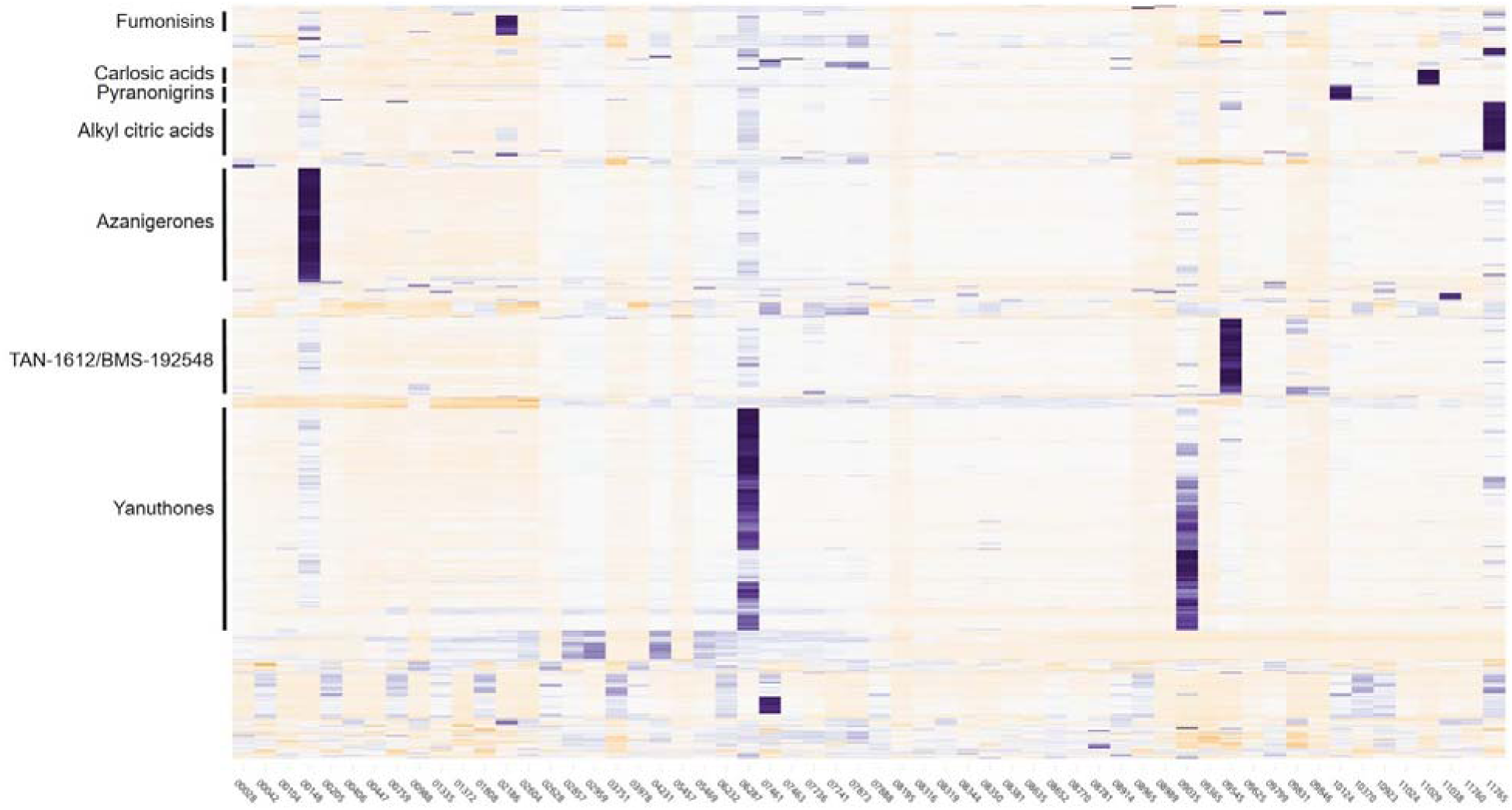
Hierarchical clustering of the secreted secondary metabolites of the transcription factor overexpressing strains. Supernatants from a five-day standing liquid culture were analysed by mass spectrometry. X-axis display the 58 strains, Y-axis display the compounds identified by FT-MS. The hierarchical clustering was generated using the statistical analysis software R (1) and the d3heatmap package (2). Dendrograms and order were obtained by calculating the Euclidian distance between samples and performing agglomerative hierarchical clustering as per the defaults of the d3heatmap package. Intensities are centered and scaled by row (across samples).

In summary, 14 of the 58 strains (∼26%) included within the overexpression strain library displayed phenotypic changes when grown on transformation or induction media. The distinct phenotypes associated with overexpression of BGC-associated TF provided initial support for our strategy and for detailed analysis of the metabolomic repertoire in our strain collection.

### Surveying the secondary metabolism of TF overexpression strains

To investigate whether overexpression of a given TF resulted in changes in the profile of secondary metabolites produced in *A. niger*, we analyzed the extracellular metabolite fraction using liquid chromatography-mass spectrometry (LC-MS). We chose to focus on extracellular metabolite production due to the propensity for fungal secondary metabolites to be secreted and to simplify the parallel analysis of metabolite production by all 58 TF overexpression strains. All TF overexpression strains were cultured in liquid medium under inducing conditions (1% or 15% maltose as a carbon source) for five or seven days. The culture supernatant was then collected, secreted metabolites were extracted and analyzed via LC-MS. The extracellular metabolite profile of each TF overexpression strain was then compared to that of the parental strain grown under the same conditions. The MS features corresponding to compounds that displayed a minimum five-fold increase in relative abundance across two biological replicates were selected for further analysis. We also analyzed the LC-MS profiles of the TF overexpression strains for production of specific metabolites using a manually curated *A. niger* metabolite database (Figure 3).

Overall, our metabolomics analysis detected the production of unique metabolites enriched more than 500-fold compared to the parental strain in 15 of 58 overexpression strains. In an additional 27 strains we identified unique metabolomic features enriched 100-fold compared to the parental strain. In the remaining 17 strains, no significant enrichment of specific metabolites was observed, or the enrichment was inconsistent across biological replicates (Figure 3). Of the 43 TF overexpression strains identified as producing unique metabolite profiles, only eight are associated with previously characterized BGCs.

Specific examples of metabolite production include overexpression of the TF NRRL3_00148. In a previous study, this TF (annotated as *azaR*) was overexpressed using a plasmid-encoded gene under control of the glyceraldehyde-3-phosphate dehydrogenase A promoter (P*_gpdA_*) (27). This led to overexpression of the cognate BGC resulting in production of several azaphilone compounds. Analysis of the strain overexpressing NRRL3_00148 in our strain collection showed a 1200-fold increase in relative abundance of a specific peak corresponding to azanigerone D (mw = 361.4) and a 500-fold increase in a peak corresponding to azanigerone E (mw = 250.2). In another example, we observed similarly significant increases in relative abundance of specific metabolites produced by the strain overexpressing the NRRL3_09545 TF. This TF has been previously shown to regulate the adjacent BGC linked to production of the TAN-1612 and BMS-192548 compounds (28). The NRRL3_09545 overexpression strain produced a 1000-fold increase in relative abundance of a compound with a *m/z* value expected of BMS-192548 (*m/z* = 414.1) (Figure 3). Taken together, these specific examples demonstrate that that stable integration of TF genes is an effective approach for eliciting the overproduction of specific secondary metabolites. By utilizing genomic integration, the strains within our collection do not require nutritional or antimicrobial selection during inductive growth, allowing for increased flexibility in potential downstream applications.

Hierarchical clustering of LC-MS data showed multifold increases in relative abundance of metabolites that could be linked to overexpression of a specific TF gene in 43 of the strains within our collection. Examples included fumonisin B_2_ that was linked to overexpression of NRRL3_02186, yanuthone D linked to overexpression of NRRL3_06287, carlosic acid and agglomerin linked to NRRL3_11029 overexpression, and hexylitaconic acid which was found in increased relative abundance in response to NRRL3_11765 overexpression. In other examples, the specific metabolites overproduced lacked matches within existing LC-MS compound databases, suggesting the production of novel compounds.

### Transcriptomic analysis of metabolite-producing TF overexpression strains

The metabolomics-guided survey of overexpression strains within our collection identified significant changes in metabolite profiles in response to TF overexpression in 43 out of 58 strains. Accordingly, we further investigated the changes in transcriptome profile in this subset of strains using RNA sequencing (RNA-seq). Strains were cultured for 24-hours under non-inducing conditions (fructose as a carbon source), then transferred to induction medium (1% maltose as carbon source) for two hours after which samples were harvested for RNA extraction. RNA-seq was performed on each sample and the data was processed into transcript-per-million (TPM) format.

Results from the RNA-seq analysis also showed strong upregulation of the specific TF genes in response to induction with 1% maltose compared to their expression in the parental strain grown under the same conditions. However, we also observed significant variation in absolute expression levels of different TF genes and fold modulation compared to the parental strain despite their identical site of insertion (glucoamylase gene locus). For example, the NRRL3_00447 TF-overexpression strain exhibited a 56-fold increase in expression of the corresponding TF gene, while in the NRRL3_08350 TF-overexpression strain the expression level of the TF gene increased 6000-fold relative to the parental NRRL2270 strain (Table S2, Figure S3).This large disparity in the magnitude of the changes in expression level occurred despite NRRL3_00447 and NRRL3_08350 showing similar expression levels, less than a 2-fold difference, in the parental strain (3.35 and 1.84 tpm, respectively).

Next, we investigated the expression level of genes residing within the cognate BGCs of the TFs targeted for overexpression. This analysis revealed three distinct scenarios. In the first scenario, we observed upregulation of expression of all genes predicted to belong to a BGC in response to TF overexpression. This occurred in nine of the overexpression strains tested. Within this set of nine strains, six represent BGCs that have been previously experimentally characterized in *A. niger*. The remaining three strains are as-yet uncharacterized experimentally and the specific chemical structures of the secondary metabolites produced in response to overexpression of these BGCs (NRRL3_04231, NRRL3_07741, NRRL3_10370) are unknown. In the second scenario we observed partial upregulation of the associated BGC, where only a subset of genes was upregulated. This occurred in six of the strains analyzed via RNA-seq (Table S2). The specific pattern of co-localized gene upregulation differed among the six strains within this category, with two distinct trends emerging: 1) backbone genes upregulated; 2) backbone genes not upregulated. More specifically, overexpression of NRRL3_02186 resulted in upregulation of 10/12 genes within the BGC. The two genes that were not upregulated include NRRL3_02187 encoding for a predicted cytochrome P450-type enzyme, and NRRL3_02188 which encodes for a predicted ABC transporter. In contrast, while overexpression of NRRL3_07461 resulted in multiple co-localized genes being upregulated, the PKS-encoding gene within this cluster (NRRL3_07443) was not upregulated.

The final scenario observed was where there was no evidence of associated BGC upregulation in response to TF overexpression. We observed this trend in the remaining 27 strains we analyzed via RNA-seq. Taken together, our transcriptional analysis suggested that overexpression of individual TF resulted in full or partial upregulation of only 26% of associated BGCs, hinting at additional layers of regulation governing BGC expression in *A. niger*.

### Upregulation of BGC correlates with overproduction of secondary metabolites

Next, we compared the LC-MS metabolomics data and the transcriptomic results for the 9 strains where we observed upregulation of the complete BGC in response to TF overexpression. Unsurprisingly, all 9 strains showed clear overproduction of secondary metabolites. Notably, this group included six strains overexpressing TFs associated with previously experimentally characterized BGCs. Overexpression of the TF (NRRL3_06287) co-localized with the BGC involved in yanuthone biosynthesis resulted in strong upregulation of the entire BGC and we observed a 53-fold enrichment in a compound with a mass corresponding to yanuthone D (mw = 502.25). The NRRL3_11765 overexpression strain also showed complete upregulation of its cognate BGC and produced a metabolite profile with multiple compounds significantly enriched compared to the parental strain. Two of the most prominent features within the metabolite profile of this strain corresponded to hexylcitrate (mw = 276.12) and hexylitatonic acid (mw = 230.11), both of which represent expected metabolic products of this BCG (Figure S4) (33). Additional overexpression strains linked to previously characterized clusters where we observed upregulation of the complete BGC and strong changes in metabolite production include those discussed above. Specifically, NRRL3_00148, NRRL3_09545, NRRL3_10124, and NRRL3_11029, all of which displayed distinct phenotypic markers in addition to robust changes in metabolite profile and complete BGC upregulation. Taken together, this result supports phenotypic changes as a strong, though imperfect (discussed below), proxy for alterations in fungal secondary metabolism.

The TFs overexpressed in the remaining three strains in this category are all associated with previously uncharacterized BGCs. In most cases, we were unable to match the additional chemical species produced by these strains with known *A. niger* metabolites, raising the possibility that they represent novel secondary metabolites. Notably, we observed overproduction of two distinct compounds in the NRRL3_07741 overexpression strain. These compounds were enriched 80-fold and 160-fold compared to the parental strain, and had molecular weights of 435.18 and 217.59, respectively. The backbone enzyme within the cluster associated with NRRL3_07741 is a NRPS and the molecular weight of these chemical species puts them within the range (200 to 3000 Da) of most known nonribosomal peptides (38). Another uncharacterized BGC showing complete gene upregulation in response to TF overexpression was associated with the TF NRRL3_10370. This strain produced a metabolite profile showing multiple compounds enriched compared to the parental strain; however, like the situation described above we were unable to identify them upon comparison to a *A. niger* metabolite database. The final uncharacterized cluster showing complete BGC upregulation also produced distinct metabolomic features. Upon overexpression of the TF NRRL3_04231, we observed 50-fold enrichment in production of tensidol B compared to the parental strain. Tensidol B is a potentiator of miconazole activity against the human pathogen *Candida albicans* (39) and had been previously isolated from the *A. niger* FKI-2342 strain; however, its biosynthetic origin is unknown. This prompted a further investigation into the possibility that this gene cluster is responsible for tensidol biosynthesis. Tensidol A and Tensidol B have been previously identified using metabolomics within extracts produced by two other *Aspergilli*, namely *A. brasiliensis* (40) *and A. tubingensis* (41, 42). A BLASTP search of genes within the BGC associated with NRRL3_04231 identified clear orthologs in both of these species, indicating a high level of synteny between *A. niger*, *A. brasiliensis,* and *A. tubingensis* at the site of this BGC (Figure 4, Table S3). A previous study that identified tensidol A and B in the metabolite extracts of *A. niger* and *A. tubingensis* found that these compounds were lacking in extracts produced by 15 other *Aspergilli* analyzed (42). Correspondingly, we initiated a search for orthologs of the NRRL3_04231-associated BGC within these species (*A. acidus*, *A. aculeatinus, A. aculeatus, A. carbonarius, A. costaricaensis, A. ellipticus, A. heteromorphus, A. homomorphus, A. ibericus, A. japonicus, A. sclerotiicarbonarius, A. sclerotiniger, A. uvarum, A. vadensis*) and found no matches in any of them. Taken together, this comparative genomic analysis provides corroborating support that the BGC containing NRRL3_04231 may be involved in tensidol biosynthesis.

**Figure 4.**
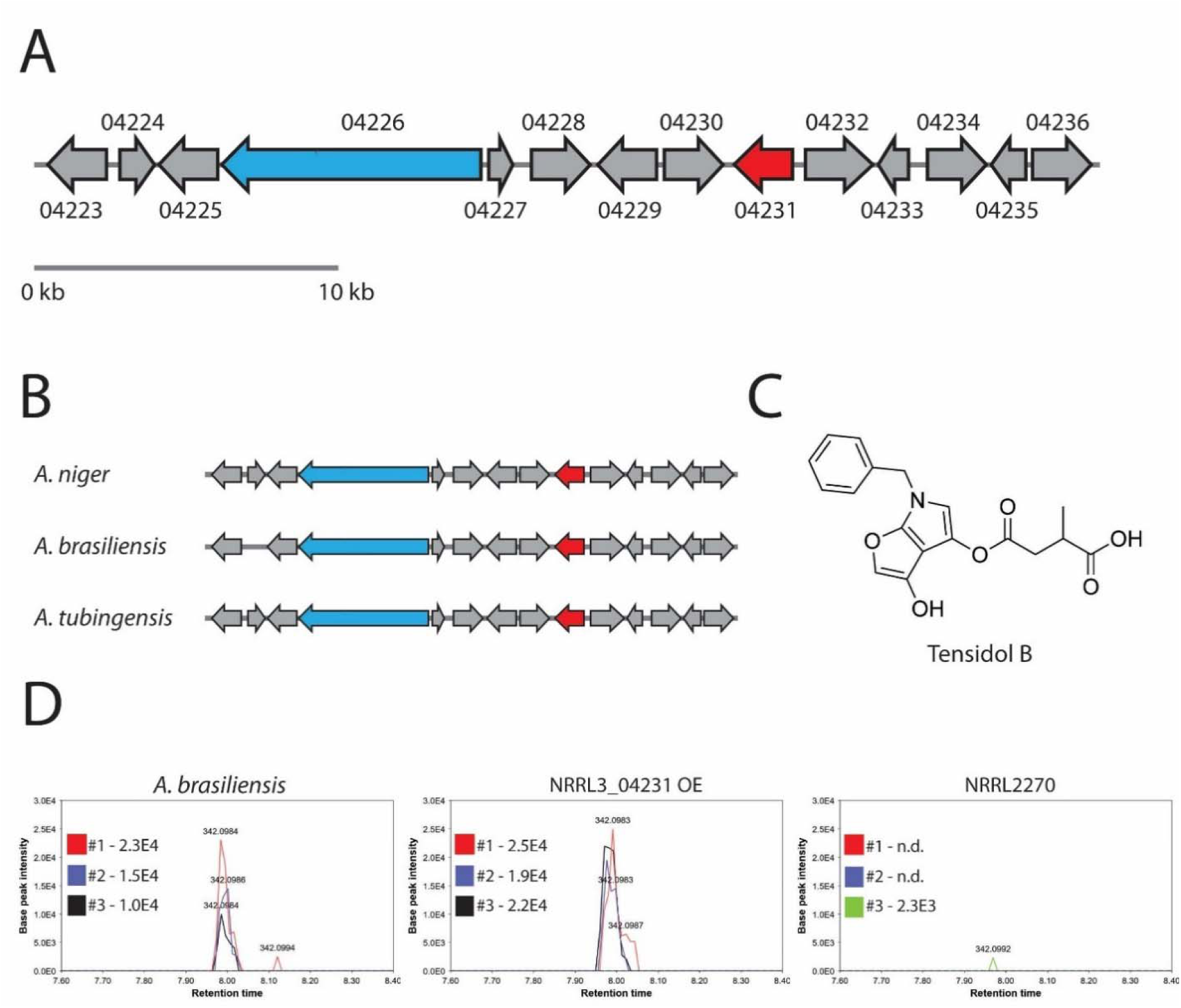
Comparative genomics implicated NRRL3_04231 in tensidol B biosynthesis. **A)** Schematic representation of the NRRL3_04231 BGC. **B)** High degree of synteny exists between *A. niger, A. brasiliensis,* and *A. tubingensis* at the site of this BGC, which is absent in other *Aspergilli* identified as non-producers of tensidols. **C)** Chemical structure of tensidol B. D) Extracted ion chromatograms (EIC) of tensidol B production found in culture extract from *A. brasiliensis*, NRRL3_04231 overexpression strain, and the parental strain NRRL2270. The peak intensity for each biological replicate is shown inset in the respective EIC panels. n.d. – not detected.

To further assess the ability of the NRRL3_04231 OE strain to produce tensidol B, we obtained a known producer of tensidol B, *A. brasiliensis* strain SN26 (ATCC 9642). We grew *A. brasiliensis* SN26 and analyzed culture extract for SM production and observed tensidol B production (*m/z* = 342.0986, rt = 7.98), which was consistent across three biological replicates. During this assessment, we also re-analyzed the NRRL3_04231 OE strain, as well as the parental strain NRRL2270 by culturing them under the same conditions (1% maltose minimal medium) and analyzing their respective culture extracts for SM production. Our results showed clear evidence of tensidol B production by the NRRL3_04231 OE strain, with a peak produced consistently across three biological replicates matching the *m/z* value and retention time observed for tensidol B in the *A. brasiliensis* culture extract. In contrast, we saw no evidence of tensidol B production within the culture extract obtained from the parental strain (Figure 4). Taken together, this result shows a clear correlation between overexpression of NRRL3_04231 and tensidol B production, providing additional experimental support implicating this cluster in tensidol biosynthesis.

### Investigating the effect of BGC partial upregulation

Further analysis of the overexpression strains with only part of the BGC upregulated by overexpression of an associated TR allowed us to investigate the specific role played by transporters encoded as part of the BGC. In two such strains, the genes encoding for predicted transporters showed little-to-no upregulation compared to the parental strain while backbone and decorating enzyme encoding genes were upregulated (Table S2). In the NRRL3_02186 overexpression strain, the corresponding BGC was predicted to produce compounds from the fumonisin class of mycotoxins based on shared similarity with fumonisin biosynthetic genes from *Fusarium verticillioides* and *Gibberella moniliformis* (24). Within this BGC, the co-localized transporter NRRL3_02188 was not upregulated in response to TF overexpression. While our LC-MS analysis of the extracellular metabolites produced by this strain detected compounds corresponding to fumonisin B1, B2, B3, B4, B5 and B6 based on matching *m/z* values and retention time scores, we hypothesized that the low expression level of the co-localized transporter could be resulting in intracellular accumulation of the secondary metabolites produced by this BGC. To test this, we extracted and analyzed the intracellular metabolites produced by the NRRL3_02186 overexpression strain and compared the relative abundance of fumonisin B2 in the intracellular and extracellular metabolite fractions. Our results showed two-fold enrichment of fumonisin B2 in the intracellular faction compared to the secreted fraction, in line with our hypothesis that low expression levels of the co-localized transporter (NRRL3_02188) was contributing to intracellular metabolite accumulation.

The second instance where we observed no upregulation of a co-localized transporter was in the NRRL3_01335 overexpression strain. This BGC is uncharacterized and its secondary metabolite products are unknown. Transcriptomic analysis revealed upregulation of 8/9 genes within the BGC, with the lone exception being a putative ABC transporter encoded by NRRL3_01330. The extracellular metabolome produced by this strain displayed modest changes compared to the parental stain. The failure of TF overexpression to elicit upregulation of the clustered ABC transporter provides a strong rationale for why we did not observe more substantive changes in metabolite production, as secondary metabolite products of this BGC may have been retained intracellularly similar to what was observed with the fumonisin BGC.

Other instances of partial upregulation were observed in the NRRL3_00104, NRRL3_06232, NRRL3_07461, and NRRL3_08989 overexpression strains. In each case, overexpression of the TF failed to upregulate the corresponding backbone enzyme(s) within these BCGs. The extracellular metabolite profile of these strains lacked unambiguous signals of additional secondary metabolite production; an observation consistent with failure to activate biosynthetic enzyme expression.

### BGC TFs are involved in a complex regulatory network

In 28 of the 43 strains where overexpression of a TF failed to upregulate its co-localized BGC we nonetheless observed evidence of additional secondary metabolite production consistent across biological replicates. This seeming discordance between the metabolomics and transcriptomics results in the case of these 28 strains prompted additional analyses of the transcriptomic data, where we identified examples that hint at further complexity and dynamism in the regulation of secondary metabolism. In the NRRL3_09035 overexpression strain we identified multiple metabolomic features enriched >500-fold compared to the parental strain (Figure 3). This set of overproduced compounds included members of the previously characterized yanuthone class of metabolites, as well as compounds for which no match was found in our existing *A. niger* compound database. The relative complexity of the metabolite profile produced in response to overexpression of NRRL3_09035 led us to hypothesize that this TF may activating BGCs in *trans*. Analysis of the transcriptome-level changes in response to overexpression of NRRL3_09035 supported this hypothesis, as multiple genes within the yanuthone BGC showed significant upregulation (Figure 5). Specifically, overexpression of NRRL3_09035 resulted in an 81-fold upregulation of the gene encoding the yanuthone BGC backbone enzyme YanA (NRRL3_06291), and 152-fold upregulation of the YanG-encoding gene, NRRL3_06290. Other genes within the yanuthone BGC were similarly upregulated, including the co-localized TF YanR (NRRL3_06287), suggesting overall activation of the yanuthone BGC in response to NRRL3_09035 overexpression may be mediated through specific upregulation of YanR (Figure 5). In this example, it appears that YanR (NRRL3_06287) may itself be under regulation by the out-of-cluster TF, NRRL3_09035.

**Figure 5.**
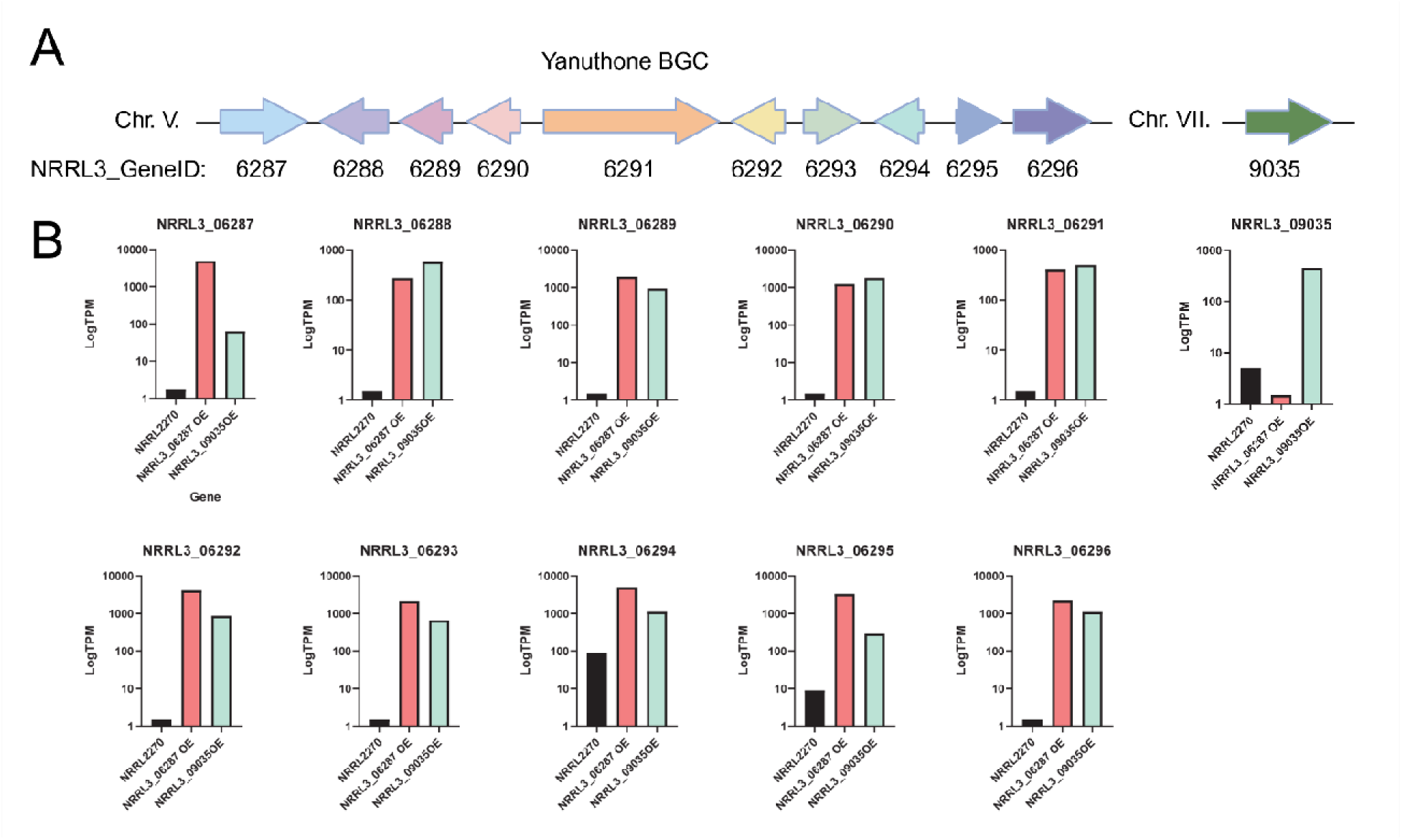
Trans activation of yanuthone BGC by NRRL3_09035. **A)** Schematic representation of the yanuthone BGC located on chromosome V. The TF NRRL3_09035 is encoded on chromosome VII. **B)** LogTPM of genes in the yanuthone BGC. Most clustered genes are silent (not expressed) in the parental strain but are overexpressed in both the NRRL3_06287 and NRRL3_09035 overexpression strains.

## Discussion

With *A. niger* serving as an industrial host for the production of proteins and organic acids, there exists a wealth of information surrounding its large-scale cultivation. This makes *A. niger* compelling for the study of secondary metabolism; however, limited understanding of control mechanisms governing the expression of fungal BGCs has long posed a challenge to uncovering, producing, and characterizing novel secondary metabolites, which has broad applications in pharma and bioindustry. As a result, the number of characterized BGCs and corresponding SM products is a small subset of the overall biosynthetic potential encoded by fungal genomes. This is readily apparent in *A. niger*, where more than 75% of its BGCs remain cryptic and uncharacterized (43). In this study, we generated a collection of *A. niger* TF overexpression strains by targeting TF genes associated with cryptic BGCs for inducible overexpression. Our subsequent analysis of these strains for phenotypic markers and changes in secreted metabolomes indicated a strong correspondence between TF overexpression and SM production; however, our transcriptomic analysis highlighted the complex and dynamic nature of BGC expression regulation.

The pipeline used for overexpression strain analysis (Figure 1) showed that phenotypic markers were a strong indicator of metabolomics changes but an imperfect proxy for gene expression. This was most apparent in our NRRL3_00205 overexpression strain, where a clear phenotype of brown pigmentation was not associated with any BGC upregulation in our transcriptomic data. We also observed metabolomics changes in many overexpression strains that did not display any obvious phenotype through the period of observation, highlighting clear practical limitations with using phenotype as an indicator of SM production.

Overall, we registered significant changes in the secreted metabolome in response to individual TR overexpression in 43 out of 58 strains produced in this study. Most of the changes corresponded to peaks and compounds that could not be reliably matched to known entities populating our *A. niger* compound database. These compounds are candidates for follow up experimentation and structural characterization. Additional assays, such as bioactivity screening, could be added to the end of our analysis pipeline to aid in prioritizing which of the candidate strains/compounds to pursue for structural characterization.

Our strategy of utilizing TF overexpression to promote SM production was based on the canonical mechanism of BGC regulation in fungal species, whereby a co-localized TF is responsible for activating gene expression of clustered biosynthetic genes. Our results provided an abundance of data that appears counter to this paradigm, and hints at regulatory complexity within *A. niger*. Our RNA-seq analysis suggested that in the majority of the overexpression strains, TF overexpression did not trigger upregulation of the adjacent BGC. This was despite them being selected for transcriptomic analysis due the presence of unique metabolomic features identified in our LC-MS analysis. In some cases, we were able to reconcile these results by identifying other non-adjacent BGCs activated in response to overexpression of a TF. The most pronounced example we observed was in the NRRL3_09035 overexpression strain, which was selected for transcriptomic analysis on account of its strong yellow pigmented phenotype and LC-MS profile. Overexpression of NRRL3_09035 failed to upregulate any of its co-localized biosynthetic genes; however, it resulted in strong upregulation of genes within the yanuthone BGC which appeared to be mediated through activation of the yanuthone TF, *yanR*. This result is a clear example or *trans*-activation of BGCs by a distant TF, and lends further credence to emerging hypotheses that there has been an over interpretation of the significance of physical clustering, suggesting a more complex relationship between TFs and associated BGCs (43). One explanation often provided for the observation of physical clustering of SM biosynthetic genes in fungal genomes is that it facilitates horizontal gene transfer of BGCs. Given this, and the multiple modalities of BGC expression (e.g., complete upregulation, partial upregulation) we observed in response to overexpression of specific TF, a possibility emerges that BGCs that adopt the more canonical mechanism of regulation (i.e., activated by a co-localized TF) may have been more recently acquired. By the same rational the BGCs that have had more time to co-evolve with their fungal host may then come under control of the more complex regulatory mechanisms we observed in the majority of our TF overexpression strains.

Additional and well-characterized parameters influencing secondary metabolism in *A. niger* are also likely to have played a role in BGC activation within our overexpression strains. Master regulatory proteins, such as McrA and LaeA, have been previously shown to influence gene expression over multiple BGCs. McrA functions as a repressor protein and has been linked to more than ten BGCs (13). A comparison between LaeA overexpression and deletion strains revealed differential expression of more than half (34 of 61 in strain FGSC A1279) of BGC-associated TFs in *Aspergillus niger*, highlighting the interplay between this methyltransferase and cluster-specific regulatory elements (44). The role of epigenetics has also been well established as a determinant in the tractability of activating BGCs (45). Previous studies have shown that treatment with small molecules known to modulate the epigenetic state of chromosomes led to changes in secondary metabolism and facilitated the discovery of new compounds (46). In cases where strains within our collection failed to display changes in gene expression and metabolite profiles, it is possible that the epigenetic state of the chromosomal location housing the BGC may have been inaccessible to the overexpressed TF. In the context of these additional layers of BGC regulation, it is possible that overexpression of individual TFs is required but insufficient to activate BGC gene expression. Consequently, it may be justified to incorporate adjunctive approaches in the future that, in addition to TF overexpression, may lead to additional strategies for efficiently activating BGC gene expression and facilitating novel small molecule discovery.

## Materials and Methods

### Manual curation of BGCs

Biosynthetic gene clusters in *A. niger* NRRL3 were manually curated following the using the publicly available NRRL3 genome sequence (available at https://pub.fungalgenomics.ca/) (47). In brief, backbone enzymes were initially identified via BLASTP searches using experimentally verified backbone enzyme sequences from other *Aspergilli* as queries (E-value ≥ 1x 10^-5^). All hits were evaluated for acceptance into their respective categories by protein domain content using the online applications Pfam (http://pfam.xfam.org/), conserved domain database (CDD) (http://www.ncbi.nlm.nih.gov/cdd/), and InterProScan 5 (http://www.ebi.ac.uk/interpro/search/sequence-search). BGCs were defined using automated tools SMURF (20) and antiSMASH (21) following the methods described in Inglis *et al*. (48). These clusters were modified to incorporate our updated gene annotation data and took precedents over the original annotations including those defined by synteny.

### Cloning of TF donor plasmids

TF genes were PCR-amplified from *A. niger* NRRL2270 genomic DNA using Phusion DNA polymerase (New England Biolabs) and the resulting PCR products were purified via a PCR Clean and Concentrator kit (Zymo). Cloning of TF genes into the pJET, a plasmid containing 655 and 700 bp of the glucoamylase promoter and the terminator regions, respectively (35), was done via ligation-independent cloning (LIC) (49). Plasmids were transformed into *E. coli* DH5_α_, purified using the Presto Mini Plasmid Kit (Geneaid) and verified via DNA sequencing. A list of primers used in this study can be found in Table S1.

### Host transformation

All TF^OE^ strains are descendants of CSFG_7003 (NRRL2270 Δ*pyrG* Δ*kusA*), a uridine auxotroph and NHEJ deficient strain that allows for high efficiency gene replacement (17). Protoplasts were prepared by incubating mycelium for three hours at 37 °C in digestion solution [40 mg/ml VinoTaste Pro (Novozymes), 1.33M sorbitol, 20 mM MES pH 5.8, 50 mM CaCl_2_]. PEG-mediated transformation was performed mostly as described in (50); briefly, protoplasts were incubated with 1 µg of ANEp8-Cas9-gRNA*glaA* and 10 µg of pJET:TF donor plasmid and plated on selective media. Two to three colonies from each transformation plate were selected and propagated on minimal media. Genomic DNA was extracted and the *glaA* locus of each candidate colony was assayed by PCR amplification using primers Fw_TF-insertion screen and Rv_TF-insertion screen. The PCR amplicon was then subjected to *Bgl*II digestion (Figure S1) to determine if gene replacement was successful based on the corresponding banding pattern.

### Growth conditions

Validated TF overexpression strains were cultured in 200 µl of either minimal media (MM) with 1% maltose or 15% maltose in 96-well plates. Plates were kept stationary and incubated at 30°C for five days.

### Sample preparation for LC-MS analysis

From standing cultures, 75 µL of culture media were collected in 1.5 mL microfuge tubes and centrifuged at 16,000 ×g for 45 min to remove mycelia, spores and cellular debris. The supernatants were transferred to new tubes and an equal volume of cold methanol (−20°C) was added for protein precipitation. Following incubation on ice for 10 min, samples were centrifuged at 16,000 × g for 45 min to remove precipitated proteins. Supernatants were transferred to fresh tubes and an equal volume of 0.1% formic acid was added. The soluble fraction was diluted 20-fold with cold 50% methanol and incubated on ice for 15 minutes with frequent mixing for protein precipitation and metabolite extraction. The samples were centrifuged for 45 min at 10 000 x g, 4°C to remove precipitated proteins. Methanol extracted metabolites were stored at -80°C until LC-MS analysis was performed.

### High-performance liquid chromatography-mass spectrometry (HPLC-MS) analysis of metabolites

HPLC-MS was performed as described in (35); briefly, 10 µL of sample was injected into a Kinetex C18 column coupled to an Agilent 1260 Infinity II HPLC System. Column flowthrough was delivered to a LTQ-FT mass spectrometer for electrospray ionization-MS. Ionization voltage was 4900 V in positive mode and 3700 V in negative mode and the scan range was 100 to 1400 m/z at a resolution of 200 m/z. Data analysis was conducted using Compound Discoverer 3.1 software and annotation was done using a custom database of 968 known *A. niger* metabolites.

### RNA-sequencing samples and analysis of data

The whole genome expression of *Aspergillus niger* parental and mutant strains was analysed using a two-step culture, a primary culture generating fungal biomass and a second transfer-culture triggering gene induction. Fifty milliliters of *Aspergillus* complete medium, 2% fructose were inoculated in 500-mL flasks with 2 x 10^6^ spores/mL and incubated 16-18 hours at 30°C, 250 rpm. From the primary cultures, wet mycelium was collected, filtered on a Buchner funnel with Miracloth (Calbiochem) and rinsed 2 times with *Aspergillus* minimum medium (51). Five milliliters (equivalent 2 grams wet weight) mycelia were transferred to 50 mL *Aspergillus* minimum medium, 1% maltose. The secondary cultures were incubated two hours at 30°C, 250 rpm. The Mycelium was collected, filtered on a Buchner funnel with Miracloth (Calbiochem), press dried with paper, flash frozen in liquid nitrogen and stored at -80°C.

Frozen mycelium was ground with a mortar and a pestle into a fine powder. Total RNA was extracted as described (52) and quantified using a NanoDrop Spectrophotometer ND-1000 (NanoDrop Technologies, Inc.) and its integrity was assessed on a 2100 Bioanalyzer (Agilent Technologies). Samples were sent to the Genome Quebec Innovation Centre for analysis; briefly, samples were loaded on a Illumina cBot and the flowcell was ran on a HiSeq 4000 for 2×100 cycles (paired-end mode). A phiX library was used as a control and mixed with libraries at 1% level. The Illumina control software was HCS HD 3.4.0.38, the real-time analysis program was RTA v. 2.7.7. Program bcl2fastq2 v2.20 was then used to demultiplex samples and generate fastq reads. RNA-sequencing data has been deposited in the National Center for Biotechnology Information database (NCBI Accession No. PRJNA1105038).

## Author Contributions

Conceptualization, A.S, A.T.; Methodology, I.B.-G., C.S.; A.T.; Validation, I.B.-G., A.S., A.T.; Investigation, Z.B., G.E., M.C.M., J-P. O., C.S., S.P., T.T.M.P., S.R.; Resources, I.B.-G., A.S., A.T.; Data Curation, M.D.F., N.G., I. R.; Writing-Original Draft Preparation, C.S.; Writing-Review & Editing, I.B.-G., A.S., A.T.; Supervision, I.B.-G., C.S.; Funding Acquisition, I.B.-G., A.S., A.T. All authors have read and agreed to the published version of the manuscript.

## Funding

This research was funded by the Industrial Biocatalysis Strategic Network (A.T., A.S.) and the Discovery Grant (I.B.-G.) of the Natural Sciences and Engineering Research Council of Canada.

## Notes

### Competing Interest Statement

The authors have declared no competing interest.

## References

1. L. Katz, R. H. Baltz, Natural product discovery: past, present, and future. J Ind Microbiol Biotechnol 43, 155–176 (2016).

2. D. J. Newman, G. M. Cragg, Natural Products as Sources of New Drugs over the Nearly Four Decades from 01/1981 to 09/2019. J Nat Prod 83, 770–803 (2020).

3. T. T. Bladt, J. C. Frisvad, P. B. Knudsen, T. O. Larsen, Anticancer and antifungal compounds from Aspergillus, Penicillium and other filamentous fungi. Molecules 18, 11338–11376 (2013).

4. T. Emri, L. Majoros, V. Toth, I. Pocsi, Echinocandins: production and applications. Appl Microbiol Biotechnol 97, 3267–3284 (2013).

5. S. Salazar-Cerezo, N. Martinez-Montiel, J. Garcia-Sanchez, Y. T. R. Perez, R. D. Martinez-Contreras, Gibberellin biosynthesis and metabolism: A convergent route for plants, fungi and bacteria. Microbiol Res 208, 85–98 (2018).

6. K. Kamei, A. Watanabe, Aspergillus mycotoxins and their effect on the host. Med Mycol 43 Suppl 1, S95–99 (2005).

7. T. Boruta, Uncovering the repertoire of fungal secondary metabolites: From Fleming’s laboratory to the International Space Station. Bioengineered 9, 12–16 (2018).

8. N. P. Keller, Fungal secondary metabolism: regulation, function and drug discovery. Nat Rev Microbiol 17, 167–180 (2019).

9. M. H. Medema et al., Minimum Information about a Biosynthetic Gene cluster. Nat Chem Biol 11, 625–631 (2015).

10. A. A. Brakhage, V. Schroeckh, Fungal secondary metabolites - strategies to activate silent gene clusters. Fungal Genet Biol 48, 15–22 (2011).

11. J. Wakefield, H. M. Hassan, M. Jaspars, R. Ebel, M. E. Rateb, Dual Induction of New Microbial Secondary Metabolites by Fungal Bacterial Co-cultivation. Front Microbiol 8, 1284 (2017).

12. T. Netzker et al., Microbial communication leading to the activation of silent fungal secondary metabolite gene clusters. Front Microbiol 6, 299 (2015).

13. C. E. Oakley et al., Discovery of McrA, a master regulator of Aspergillus secondary metabolism. Mol Microbiol 103, 347–365 (2017).

14. K. D. Clevenger et al., A scalable platform to identify fungal secondary metabolites and their gene clusters. Nat Chem Biol 13, 895–901 (2017).

15. C. Liu et al., In vitro reconstitution of a PKS pathway for the biosynthesis of galbonolides in Streptomyces sp. LZ35. Chembiochem 16, 998–1007 (2015).

16. T. C. Cairns, C. Nai, V. Meyer, How a fungus shapes biotechnology: 100 years of Aspergillus niger research. Fungal Biol Biotechnol 5, 13 (2018).

17. L. Song, J. P. Ouedraogo, M. Kolbusz, T. T. M. Nguyen, A. Tsang, Efficient genome editing using tRNA promoter-driven CRISPR/Cas9 gRNA in Aspergillus niger. PLoS One 13, e0202868 (2018).

18. J. Brandl, M. R. Andersen, Aspergilli: Models for systems biology in filamentous fungi. Current Opinion in Systems Biology 6, 67–73 (2017).

19. M. V. Aguilar-Pontes et al., The gold-standard genome of Aspergillus niger NRRL 3 enables a detailed view of the diversity of sugar catabolism in fungi. Stud Mycol 91, 61–78 (2018).

20. N. Khaldi et al., SMURF: Genomic mapping of fungal secondary metabolite clusters. Fungal Genet Biol 47, 736–741 (2010).

21. K. Blin et al., antiSMASH 7.0: new and improved predictions for detection, regulation, chemical structures and visualisation. Nucleic Acids Res 51, W46–W50 (2023).

22. K. Blin et al., antiSMASH 5.0: updates to the secondary metabolite genome mining pipeline. Nucleic Acids Res 47, W81–W87 (2019).

23. A. Susca et al., Variation in the fumonisin biosynthetic gene cluster in fumonisin-producing and nonproducing black aspergilli. Fungal Genet Biol 73, 39–52 (2014).

24. R. H. Proctor, D. W. Brown, R. D. Plattner, A. E. Desjardins, Co-expression of 15 contiguous genes delineates a fumonisin biosynthetic gene cluster in Gibberella moniliformis. Fungal Genet Biol 38, 237–249 (2003).

25. A. Susca et al., Variation in Fumonisin and Ochratoxin Production Associated with Differences in Biosynthetic Gene Content in Aspergillus niger and A. welwitschiae Isolates from Multiple Crop and Geographic Origins. Front Microbiol 7, 1412 (2016).

26. S. Theobald et al., Uncovering secondary metabolite evolution and biosynthesis using gene cluster networks and genetic dereplication. Sci Rep 8, 17957 (2018).

27. A. O. Zabala, W. Xu, Y. H. Chooi, Y. Tang, Characterization of a silent azaphilone gene cluster from Aspergillus niger ATCC 1015 reveals a hydroxylation-mediated pyran-ring formation. Chem Biol 19, 1049–1059 (2012).

28. Y. Li, Y. H. Chooi, Y. Sheng, J. S. Valentine, Y. Tang, Comparative characterization of fungal anthracenone and naphthacenedione biosynthetic pathways reveals an alpha-hydroxylation-dependent Claisen-like cyclization catalyzed by a dimanganese thioesterase. J Am Chem Soc 133, 15773–15785 (2011).

29. X. L. Yang, T. Awakawa, T. Wakimoto, I. Abe, Three acyltetronic acid derivatives: noncanonical cryptic polyketides from Aspergillus niger identified by genome mining. Chembiochem 15, 1578–1583 (2014).

30. S. E. Baker, Aspergillus niger genomics: past, present and into the future. Med Mycol 44 Suppl 1, S17–21 (2006).

31. D. J. Jeenes, B. Marczinke, D. A. MacKenzie, D. B. Archer, A truncated glucoamylase gene fusion for heterologous protein secretion from Aspergillus niger. FEMS Microbiol Lett 107, 267–271 (1993).

32. J. M. Withers et al., Optimization and stability of glucoamylase production by recombinant strains of Aspergillus niger in chemostat culture. Biotechnol Bioeng 59, 407–418 (1998).

33. S. Palys, T. T. M. Pham, A. Tsang, Biosynthesis of Alkylcitric Acids in Aspergillus niger Involves Both Co-localized and Unlinked Genes. Front Microbiol 11, 1378 (2020).

34. Y. M. Chiang et al., Characterization of a polyketide synthase in Aspergillus niger whose product is a precursor for both dihydroxynaphthalene (DHN) melanin and naphtho-gamma-pyrone. Fungal Genet Biol 48, 430–437 (2011).

35. G. Evdokias et al., Identification of a Novel Biosynthetic Gene Cluster in Aspergillus niger Using Comparative Genomics. J Fungi (Basel) 7 (2021).

36. D. K. Holm et al., Molecular and chemical characterization of the biosynthesis of the 6-MSA-derived meroterpenoid yanuthone D in Aspergillus niger. Chem Biol 21, 519–529 (2014).

37. T. Awakawa, X. L. Yang, T. Wakimoto, I. Abe, Pyranonigrin E: a PKS-NRPS hybrid metabolite from Aspergillus niger identified by genome mining. Chembiochem 14, 2095–2099 (2013).

38. M. Chevalier et al., Kendrick Mass Defect Approach Combined to NORINE Database for Molecular Formula Assignment of Nonribosomal Peptides. J Am Soc Mass Spectrom 30, 2608–2616 (2019).

39. T. Fukuda et al., Tensidols, new potentiators of antifungal miconazole activity, produced by Aspergillus niger FKI-2342. J Antibiot (Tokyo) 59, 480–485 (2006).

40. J. Varga et al., Aspergillus brasiliensis sp. nov., a biseriate black Aspergillus species with world-wide distribution. Int J Syst Evol Microbiol 57, 1925–1932 (2007).

41. J. Varga et al., New and revisited species in Aspergillus section Nigri. Stud Mycol 69, 1–17 (2011).

42. R. A. Samson et al., Diagnostic tools to identify black aspergilli. Stud Mycol 59, 129–145 (2007).

43. M. J. Kwon et al., Beyond the Biosynthetic Gene Cluster Paradigm: Genome-Wide Coexpression Networks Connect Clustered and Unclustered Transcription Factors to Secondary Metabolic Pathways. Microbiol Spectr 9, e0089821 (2021).

44. B. Wang et al., Profiling of secondary metabolite gene clusters regulated by LaeA in Aspergillus niger FGSC A1279 based on genome sequencing and transcriptome analysis. Res Microbiol 169, 67–77 (2018).

45. J. C. Albright et al., Large-scale metabolomics reveals a complex response of Aspergillus nidulans to epigenetic perturbation. ACS Chem Biol 10, 1535–1541 (2015).

46. K. M. Fisch et al., Chemical induction of silent biosynthetic pathway transcription in Aspergillus niger. J Ind Microbiol Biotechnol 36, 1199–1213 (2009).

47. H. Almeida, S. Palys, A. Tsang, A. B. Diallo, TOUCAN: a framework for fungal biosynthetic gene cluster discovery. NAR Genom Bioinform 2, lqaa098 (2020).

48. D. O. Inglis et al., Comprehensive annotation of secondary metabolite biosynthetic genes and gene clusters of Aspergillus nidulans, A. fumigatus, A. niger and A. oryzae. BMC Microbiol 13, 91 (2013).

49. F. Thieme, C. Engler, R. Kandzia, S. Marillonnet, Quick and clean cloning: a ligation-independent cloning strategy for selective cloning of specific PCR products from non-specific mixes. PLoS One 6, e20556 (2011).

50. V. Meyer et al., Fungal gene expression on demand: an inducible, tunable, and metabolism-independent expression system for Aspergillus niger. Appl Environ Microbiol 77, 2975–2983 (2011).

51. E. Kafer, Meiotic and mitotic recombination in Aspergillus and its chromosomal aberrations. Adv Genet 19, 33–131 (1977).

52. N. Semova et al., Generation, annotation, and analysis of an extensive Aspergillus niger EST collection. BMC Microbiol 6, 7 (2006).

53. H. F. Tsai, M. H. Wheeler, Y. C. Chang, K. J. Kwon-Chung, A developmentally regulated gene cluster involved in conidial pigment biosynthesis in Aspergillus fumigatus. J Bacteriol 181, 6469–6477 (1999).

54. C. Gil Girol et al., Regio- and stereoselective oxidative phenol coupling in Aspergillus niger. Angew Chem Int Ed Engl 51, 9788–9791 (2012).

55. Y. Lv, J. Xiao, L. Pan, Type III polyketide synthase is involved in the biosynthesis of protocatechuic acid in Aspergillus niger. Biotechnol Lett 36, 2303–2310 (2014).

